# Cerebellar structural and functional alterations during morphine self-administration are associated with motivation and disrupted goal-directed actions in male Wistar rats

**DOI:** 10.64898/2026.04.23.720465

**Authors:** Mariana Stefania Serrano-Ramírez, Jalil Rasgado-Toledo, David Medina-Sánchez, Luis A. Trujillo-Villarreal, César J. Carranza-Aguilar, Eduardo A. Garza-Villarreal

## Abstract

Opioid addiction is characterized by a strong motivational drive to obtain the drug, traditionally attributed to neuroadaptations within mesocorticolimbic reward circuits. Increasing evidence suggests that opioid-induced plasticity extends beyond these pathways, implicating the cerebellum in addiction, although its contribution to drug seeking remains poorly defined. Here, we hypothesized that morphine self-administration would induce cerebellum-associated behavioral alterations alongside structural and functional remodeling. Male Wistar rats (N = 27) were trained to self-administer morphine (0.1 mg/kg/infusion) or saline under fixed-ratio (FR1, 3 h/day, 20 days) and progressive-ratio (PR 9-4, 6 h/day, 25 days) schedules. Behavioral assessments included open field, elevated plus maze, novel object recognition, and Morris water maze tasks. Structural and functional magnetic resonance imaging was acquired longitudinally using a 7T scanner. Morphine self-administering rats showed a progressive increase in infusions and motivation. Exploratory activity increased without affecting anxiety-like behavior or recognition memory. In the Morris water maze, spatial learning was preserved; however, rats exhibited increased Whishaw’s error (path complexity) and a shift from serial to disorganized navigation strategies, indicating impaired movement sequencing. We also found cerebellar volume changes in Crus1, 7Cb, and 8Cb that were accompanied by altered cerebello-cerebral functional connectivity with insular, striatal, hippocampal, motor, and reticular networks. Multivariate analysis further suggested a brain–behavior covariance pattern in which motivational measures showed the strongest contribution, involving a distributed structural network including the cerebellum and insular cortex. These findings indicate that morphine self-administration preserves drug-seeking motivation while disrupting the organization of goal-directed actions, alongside cerebellar remodeling and altered cerebello-cerebral coupling.

## 1. Introduction

Opioid addiction is a major public health concern characterized by compulsive drug seeking, impaired control over intake, and vulnerability to relapse (Koob and Volkow 2016). Opioids such as morphine exert their pharmacological actions primarily through μ-opioid receptors, producing acute analgesia via modulation of nociceptive pathways, while their rewarding effects are mediated by mesocorticolimbic circuits (Fields 2004; Koob and Volkow 2016). With repeated exposure, these effects give rise to well-characterized neuroadaptations that support the transition from controlled drug use to compulsive seeking (Koob 2021; Carranza-Aguilar et al. 2022). While reward-centered neurobiological models account for reinforcement and motivational drive, they are insufficient to explain the full spectrum of cognitive, affective, and motor disturbances associated with chronic opioid exposure (Winstanley et al. 2021; Quinn et al. 2023; Paul et al. 2021). These observations suggest that opioid-induced plasticity extends beyond canonical reward circuits and involves additional neural systems related to the regulation, organization, and execution of behavior.

Increasing attention has been directed toward neural structures that integrate motivational signals with behavioral output. Among these, the cerebellum has emerged as a key candidate due to its integrative role across motor, cognitive, and affective domains (Moulton et al. 2014; Moreno-Rius and Miquel 2017; Rudolph et al. 2023). Beyond its classical involvement in motor coordination, the cerebellum contributes to goal-directed actions, visuospatial strategy selection, and predictive sensorimotor control (Ramnani 2012; Strick et al. 2009). Within this view, cerebellar activity increases during tasks that place demands on internal modeling and predictive control (Manto et al. 2024), underscoring its role in shaping structured behavioral responses. The relevance of cerebellar functions becomes evident in addiction-related contexts, where drug seeking depends not only on reward pursuit but also on the translation of signals into coherent, goal-directed action patterns (Moulton et al. 2014; Everitt and Robbins 2016). Neuroimaging studies in opioid-dependent humans have reported cerebellar activation during cue-induced craving and drug-related subjective states (Sell et al. 2000), and converging evidence suggests that the cerebellum directly modulates reward-related circuits through projections to the ventral tegmental area, a key region in the mesolimbic system that contribute to addiction-related behaviors (Carta et al. 2019).

Cerebellar circuits may be sensitive to opioid exposure due to the presence of μ-opioid receptors within cerebellar neurons and local circuitry (Kinney and White 1991; Bekheet et al. 2010; Yang et al. 2022). Human neuroimaging studies have documented cerebellar structural alterations in heroin-dependent individuals (Wang et al. 2013), as well as reductions in cerebellar gray matter volume in heroin users on methadone maintenance treatment (Lin et al. 2012). While these findings establish the relevance of cerebellum in opioid use disorder, animal models offer a critical advantage for mechanistic investigation, as they allow the dissociation of direct pharmacological effects from the operant components of voluntary drug taking. In rats, passive morphine administration produces cerebellar alterations, including reductions in neuronal soma size (Bekheet et al. 2010). Extending this, in our recent work using morphine self-administration we showed that cerebellar remodeling can emerge early during voluntary opioid intake, even in the absence of neuronal loss, suggesting adaptive rather than neurodegenerative processes (Elizarrarás-Herrera et al. 2026). Parallel observations of cerebellar alterations following exposure to other substances of abuse, such as cocaine and alcohol, further suggest that this region may contribute to addiction-related neurobiological processes in a manner that is not restricted to a single drug class (Vazquez-Sanroman et al. 2015; Gómez-Villatoro et al. 2025).

Characterizing cerebellar alterations produced by morphine self-administration and determining how these changes relate to behavior is critical for defining the role of this structure in addiction-related processes. To examine how morphine self-administration affects cerebellar structure, function, and related behavioral performance, we conducted a longitudinal study in male Wistar rats trained to self-administer morphine, using *in vivo* MRI to monitor cerebellar changes as drug-taking behavior progressed. The aims of the study were to: 1) characterize brain volume changes across a morphine self-administration protocol, 2) evaluate cerebellum-associated behaviors, including exploration patterns and navigation strategies, 3) assess how morphine self-administration modifies the cerebellar-cerebral functional connectivity, particularly within motivational, interoceptive, and motor-related networks, and 4) integrate structural and behavioral measures using multivariate analysis to identify brain-behavior relationships associated with morphine seeking. Through this approach, we sought to clarify the role of the cerebellum in opioid-seeking behavior.

## 2. Methods and materials

### Animals

Male Wistar rats were obtained from our institutional vivarium and individually housed in ventilated Plexiglas cages under an inverted 12:12 h light/dark cycle (lights on at 10:00 a.m.), with controlled temperature (22-24 °C) and *ad libitum* access to food and water. Animals were handled for 15 minutes per day to habituate them to manipulation. All procedures were approved by the Bioethics Committee of the Institute of Neurobiology (Protocol 113.A) and conducted in accordance with the Mexican Official Standard for the care and handling of laboratory animals (NOM-062-ZOO-1999), the Guide for the Care and Use of Laboratory Animals (National Research Council (US) 2011), and the ARRIVE 2.0 guidelines (Percie du Sert et al. 2020).

### Drugs

Morphine sulfate (Biogenfine®) was administered intravenously at a dose of 0.1 mg/kg/infusion. Isoflurane (Sofloran®) was used for anesthesia during surgery and MRI procedures (5% induction limited to a maximum duration of two minutes, ∼1 % maintenance). For functional MRI, dexmedetomidine (0.012 mg/kg, s.c.) was administered as a single bolus, 20 min before scanning. Meloxicam (0.3 mg/kg) was administered as postoperative analgesia, and enrofloxacin (10 mg/kg) as a preventive antibiotic. Catheter patency was maintained with a ticarcillin (66.7 mg/ml) and heparin (20 IU/ml) flushing solution. Patency was verified when obstruction was suspected by administering a ketamine (15 mg/ml) and midazolam (0.75 mg/ml) mixture, which induces a brief loss of the righting reflex starting within 10 seconds of injection. All drugs were obtained from Sigma-Aldrich (Toluca, Mexico).

### Experimental Design

Twenty-seven rats were initially included in this study (Fig. 1). Animals were habituated to handling from postnatal day (P) 40 to P42 (15 min/day) and then exposed to the operant chambers and a protocol of successive approximations per three days using food reinforcement. Jugular catheter surgery was performed on P46, and animals completed a six-day recovery period. Once recovered, rats were randomly assigned to the morphine (n = 11) or control (n = 9) groups, whereas 7 animals were excluded due to catheter-related complications. Morphine self-administration began on P59 under a fixed-ratio 1 schedule (FR1; 3 hours/day) and continued for twenty days, after which animals completed a twenty-five-day period under a progressive-ratio 9-4 schedule (PR 9-4; 6 hours/day). Early behavioral assessments were performed during FR1 period, including the open field on P75 and the elevated plus maze on P76. Animals then progressed through the extended behavioral battery, which comprised novel object recognition (NOR) between P87 and P89, open field test (OFT) on P90, the elevated plus maze (EPM) on P92, and the spatial Morris Water Maze (MWM) from P94 to P98. Structural MRI scans were obtained at three predefined time points with 1-2 days of scanning period: P57/58 (T1), P69/70 (T2) and P104/105 (T3).

**Figure 1.**
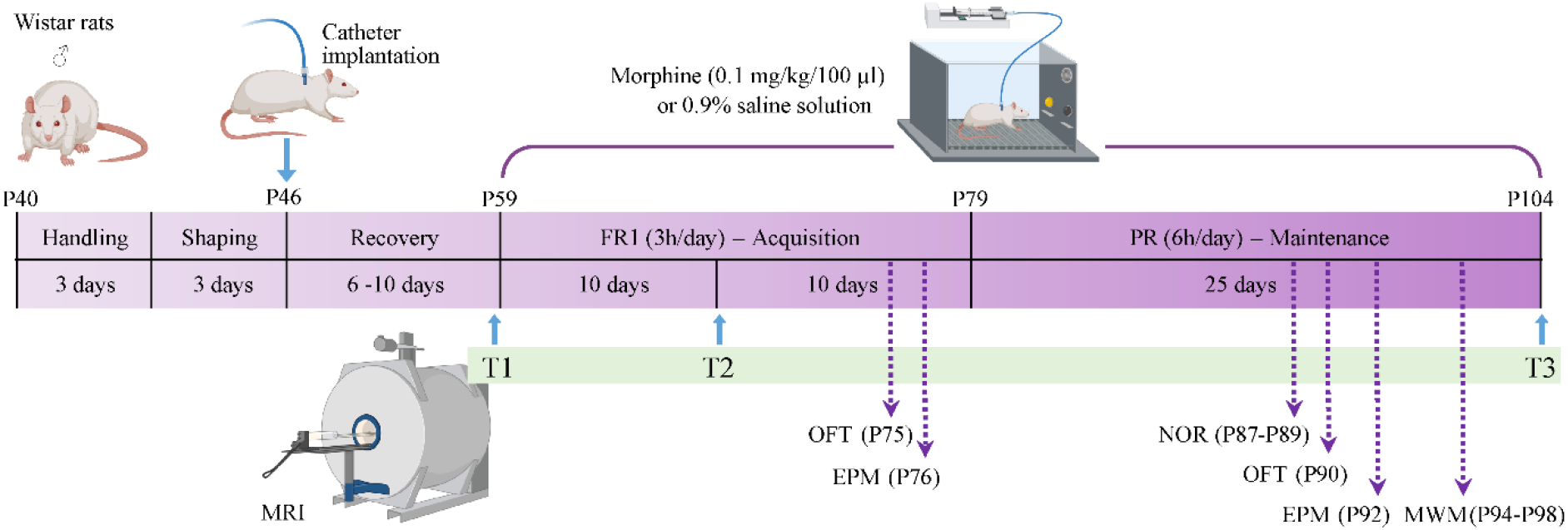
Experimental design. Male Wistar rats were obtained on postnatal day (P) 40 and habituated to handling (15 min/day) before undergoing a food-reinforced successive-approximations protocol. Jugular catheter surgery was performed on P46, followed by a recovery period. On P58, animals were assigned to morphine or saline control groups and began intravenous self-administration under a fixed-ratio 1 (FR1) schedule for 20 days (3 h/day), followed by a progressive-ratio 9-4 (PR 9-4) schedule lasting 25 days (6 h/day). Each infusion was delivered over 5 s, paired with a cue light, and followed by a 30-s timeout. Structural and resting-state functional MRI scans were acquired at three time points: baseline (P57–P58, T1), mid-training (P69–P70, T2), and after completion of the protocol (P104–P105, T3). During the FR1 phase, behavioral assessments included the open field test (P75) and elevated plus maze (P76). During the PR phase, animals completed an extended behavioral battery comprising novel object recognition (NOR; P87–P89), open field (OF; P90), elevated plus maze (EPM; P92), and the Morris water maze (MWM; P94–P98). P, Postnatal day; Mor, Morphine; Ctrl, Control; FR1, Fixed Ratio 1; PR, Progressive Ratio; MRI, Magnetic Resonance Imaging; NOR, Novel Object Recognition; OF, Open Field; EPM, Elevated Plus Maze; MWM. Partially created in BioRender. Rasgado, J. (2026) https://BioRender.com/mizeuds.

### Catheter implantation

Under isoflurane anesthesia (5% induction, 1% maintenance), a catheter was implanted into the right jugular vein, tunneled subcutaneously to the interscapular region, and connected to a non-magnetic vascular access button. Animals underwent a six-day recovery period with postoperative analgesia (meloxicam, 0.3 mg/kg) and prophylactic antibiotic treatment (enrofloxacin, 10 mg/kg). Catheter patency was maintained by daily flushing with heparinized saline containing ticarcillin (66.67 mg/ml) and heparin (20 IU/ml) and verified by a ketamine (15 mg/ml) and midazolam (0.75 mg/ml) challenge inducing rapid loss of the righting reflex. After recovery, catheters were flushed daily with heparinized saline following each self-administration session and monitored for complications.

### Operant training and self-administration schedules

Rats were trained in automated operant chambers (Med-Associates, ENV-018V) using a shaping protocol based on reinforced successive approximations (Elizarrarás-Herrera et al. 2026). Animals were then introduced to a two-lever configuration to establish drug-reinforced responding. Self-administration sessions were conducted during the dark phase. Under a fixed-ratio 1 schedule (FR1; 20 days), each active lever press delivered morphine (0.1 mg/kg/infusion), followed by a progressive-ratio 9–4 schedule (PR 9–4; 6 h/day, 25 days) with escalating response requirements (Grasing et al. 2003). Lever presses and infusions were recorded. Motivation was calculated from the final five PR sessions as the mean number of active lever presses emitted before the last infusion (Bock et al. 2013; Le et al. 2017).

### Behavioral tests

Exploratory activity was quantified in the OFT. Anxiety-like behavior was assessed in the EPM using an anxiety index, and recognition memory in the NOR using a discrimination ratio. Spatial learning and navigation were evaluated in the MWM during acquisition and probe sessions, including escape latency, distance and directionality to the former platform location, time and entries in the target quadrant, Whishaw’s error, and swimming strategy classification.

### Magnetic Resonance Imaging

Magnetic resonance imaging (MRI) was performed on a 7T Bruker Pharmascan system using ParaVision v7.0. High-resolution structural images were acquired using a T2-weighted 3D FLASH sequence (160 µm isotropic resolution). Resting-state fMRI data were obtained using a T2*-weighted gradient-echo echo-planar imaging (GE-EPI) sequence (10-min acquisition). Structural data were analyzed using longitudinal deformation-based morphometry (DBM) to estimate regional volumetric changes across time, with anatomical labeling based on the SIGMA rat brain atlas (Barrière et al., 2019). Resting-state fMRI data were preprocessed using RABIES, and seed-based functional connectivity analyses were performed using cerebellar clusters identified from the DBM results.

### Statistical analysis

Self-administration and behavioral measures were analyzed using linear mixed-effects models with rat as a random factor and group and time-related variables as fixed effects. Probe-test outcomes were evaluated using regression-based approaches, including generalized linear models for count data and contingency analyses. Structural MRI data were analyzed using voxel-wise mixed-effects models (Group × Age or Age^2^) with batch as a covariate and false discovery rate (FDR) correction. Functional connectivity differences were assessed using mixed-effects modeling. Brain-behavior associations were examined using partial least squares (PLS) correlation with permutation testing. Significance thresholds were p < 0.05 for behavioral analyses and q < 0.05 for voxel-wise results. All analyses were performed in R v4.2.3 (RRID:SCR_001905).

*Additional methodological details are provided in the Supplementary Material*.

## 3. Results

### Progressive acquisition of morphine self-administration and increased motivation

Morphine self-administration was evaluated across two operant phases. During FR1 acquisition (sessions 1-20), morphine-exposed rats progressively increased drug intake relative to controls, with significant differences emerging at sessions 7, 8, 11, 14, and 18 (Fig. 2A). During the PR phase (sessions 21-45), morphine-treated rats showed an escalation in infusions (F_(1,826)_ = 207.99, p ≤ 0.001) with significant group × session interaction (F_(43,826)_ = 2.37, p ≤ 0.0001; Fig. 2A). Correct lever presses displayed a parallel pattern, with morphine-exposed rats showing significantly more active responding at FR1 sessions 6, 7, 8, 11, and 19, and at PR sessions 21, 23-26, 29, 30, and 35 (Fig. 2B). Inactive lever presses remained low overall, with only session 27 showing a significant difference (Fig. 2C). Motivational scores further separated the groups, with saline animals clustering near the origin and morphine-self-administered rats exhibiting higher scores that scaled positively with infusion number (Fig. 2D), consistent with the emergence of a compulsive-like seeking profile.

**Figure 2.**
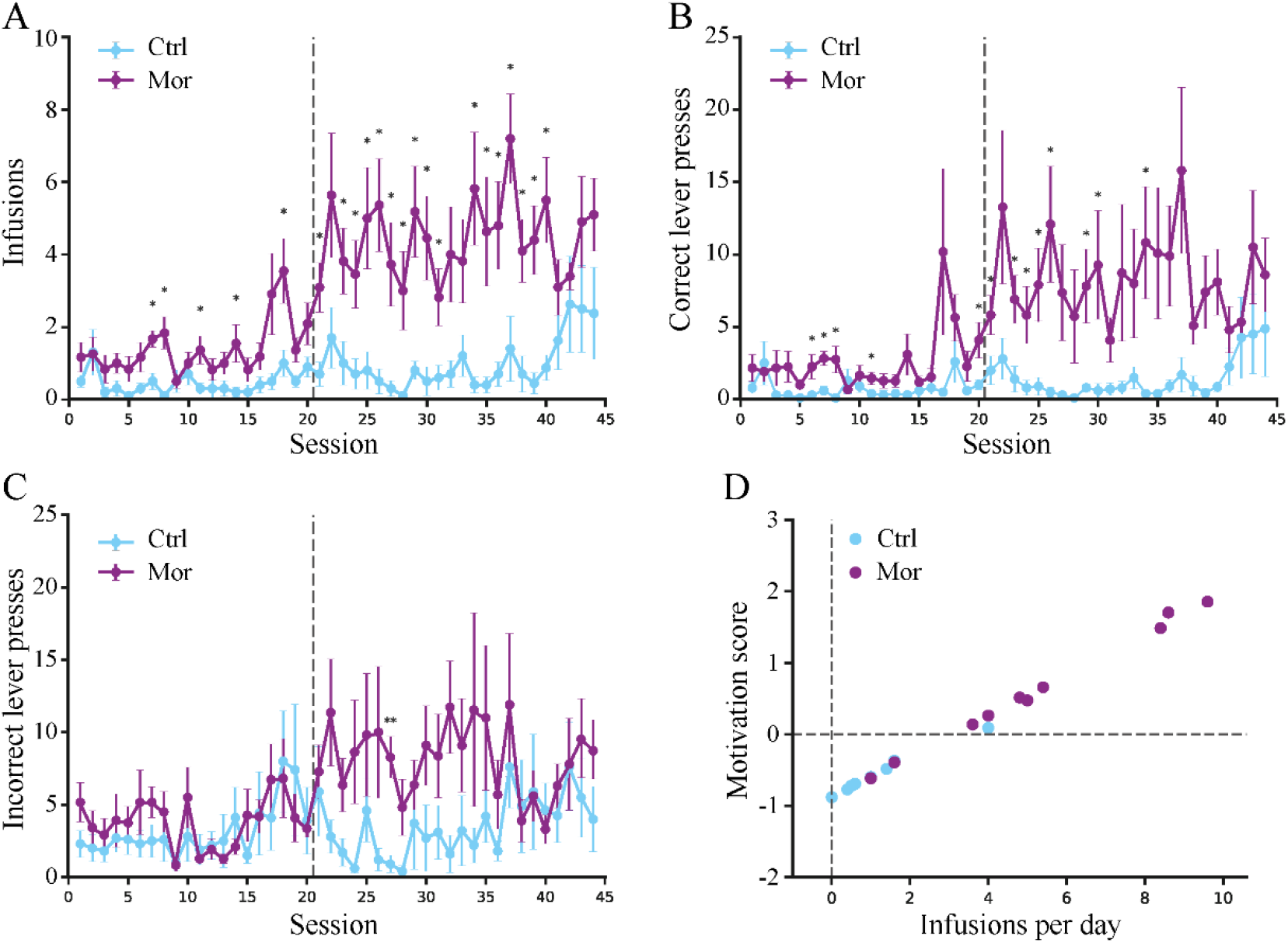
Morphine self-administration behavior and motivational performance. **A.**Number of infusions per session across operant training under a fixed-ratio 1 (FR1; sessions 1–20) and a progressive-ratio 9-4 (PR; sessions 21–44) schedule. **B**. Total correct (active) lever presses per session. **C**. Total incorrect (inactive) lever presses. The vertical dashed line indicates the transition from FR1 (1-20) to PR (21-45) schedules. **D**. Motivation score (z-scores) plotted against the average number of infusions per day for individual animals. Data are shown as group means ± standard error of the mean (SEM) for Ctrl and Mor groups. *p < 0.05, **p < 0.01, LMM, followed by Tukey-corrected post hoc comparisons.

### Morphine self-administration increases exploration and alters spatial navigation patterns

Animals were evaluated using multiple tasks (Fig. 1). In the OFT, rearing behavior differed between groups for both count (Fig. 3A; F_(1,18)_ = 7.07, p = 0.016, η^2^g = 0.22) and duration (Fig. 3B; F_(1,18)_ = 5.42, p = 0.032, η^2^g = 0.19). Group differences became evident during the second assessment (T2), when morphine intake was already established. At this point, the morphine group displayed a significantly higher number of rearing (t_(18)_ = 3.16, p = 0.005; Fig. 3A) and longer rearing episodes (t_(18)_ = 2.79, p = 0.012; Fig. 3B), indicative of enhanced exploratory behavior. In the EPM, no significant effects of group or phase, nor a group × phase interaction, were observed for anxiety index (all p > 0.43; η^2^g ≤ 0.011; Fig. 3C). In the NOR task, assessed after stabilization of drug-taking behavior, morphine-exposed rats showed a slightly lower discrimination index than controls; however, this difference did not reach statistical significance (t_(15.19)_ = 1.92, p = 0.074; g = 0.89, 95% CI [-0.08, 1.84]; Fig. 3D). Subsequently, animals were tested in the MWM to evaluate spatial learning, memory retention, and navigational precision. During acquisition, both groups showed a progressive reduction in escape latency across training days, indicating successful spatial learning (F_(3,54)_ = 13.24, p < 0.001; R^2^ = 0.33). There were no effects of group (p = 0.82) and no group × day interaction (p = 0.57; Fig. 3E), demonstrating comparable learning rates between morphine and control groups. During the probe test, both groups exhibited the expected preference for the quadrant that previously contained the platform, indicating preserved spatial memory retention. No group differences were observed across probe-test measures, including distance to the former platform location (t_(15)_ = −0.69, p = 0.751, g = −0.32), directionality toward the target (t_(15)_ = 0.32, p = 0.375, g = 0.15), time spent in the target quadrant (t_(15)_ = −0.49, p = 0.685, g = −0.23), and number of entries into that quadrant (z = 0.87, p = 0.382, IRR = 1.34; Supplementary Table S1). In contrast, morphine-exposed rats exhibited higher Whishaw’s error values than controls (t_(15)_ = 1.93, p = 0.036, g = 0.90; Fig. 3F), indicating reduced trajectory precision despite preserved spatial memory retention.

**Figure 3.**
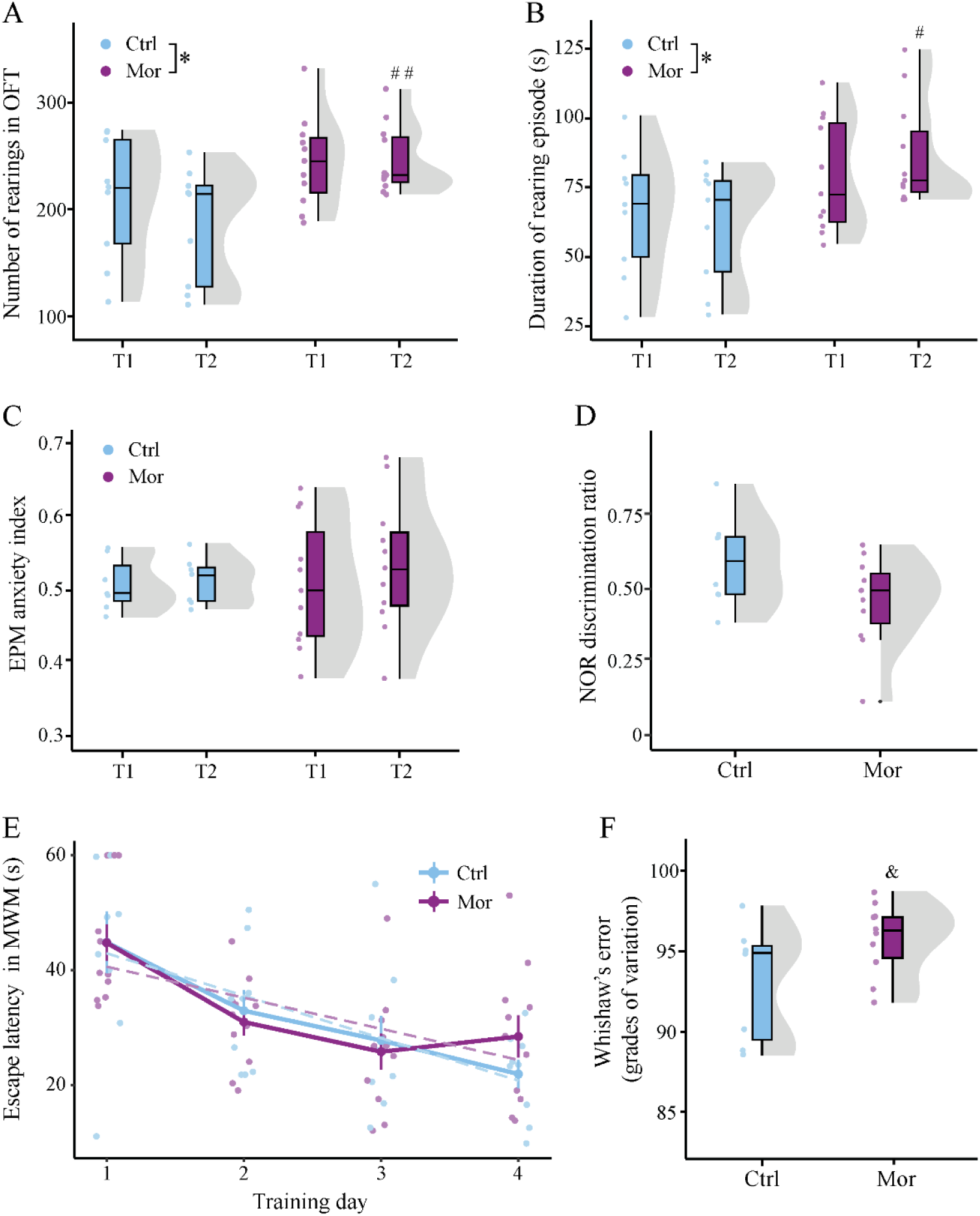
Effects of morphine self-administration on exploratory behavior, anxiety-like behavior, recognition memory, and spatial performance. **A.**Number of rearings in the Open Field Test (OFT) during the first (T1) and second (T2) assessments. **B**. Duration of rearing episodes in the OFT at T1 and T2. **C**. Anxiety index in the Elevated Plus Maze (EPM) at T1 and T2. **D**. Discrimination index in the Novel Object Recognition (NOR) test. **E**. Escape latency across training days in the Morris Water Maze (MWM). **F**. Whishaw’s error (angular deviation index of swimming trajectory precision toward the former platform location) during the MWM probe test (day 5). Data are presented as mean ± SEM. *p < 0.05, **p < 0.01, two-way repeated-measures ANOVA (Group × Phase). ^#^p < 0.05, ^##^p < 0.01, Bonferroni-adjusted post hoc comparisons. ^&^p < 0.05, ordinary least squares regression.

### Morphine exposure impairs navigation precision and abolishes serial strategies despite preserved spatial learning

To examine whether alterations in navigational precision extended to goal-directed behavior and movement organization, navigational strategies were examined. A significant group difference in the distribution of trajectory types was detected (p = 0.0368; V = 0.47; Fig. 4). In specific, the proportion of spatial trajectories was similar between groups (Mor: 50.0% vs. Ctrl: 28.6%; p = 0.622; Fig. 4A), serial trajectories were completely absent in the morphine group but were present in controls (0% vs. 57.1%; p = 0.0147; φ = 0.66; Fig. 4B), and random trajectories were more frequent in the morphine group (50.0%) than in controls (14.3%), although this difference did not reach statistical significance (p = 0.304; φ = 0.37; Fig. 4C). Overall, these results suggest that morphine self-administration altered swimming strategy selection, characterized primarily by the elimination of serial trajectories commonly associated with organized, goal-directed navigation.

**Figure 4.**
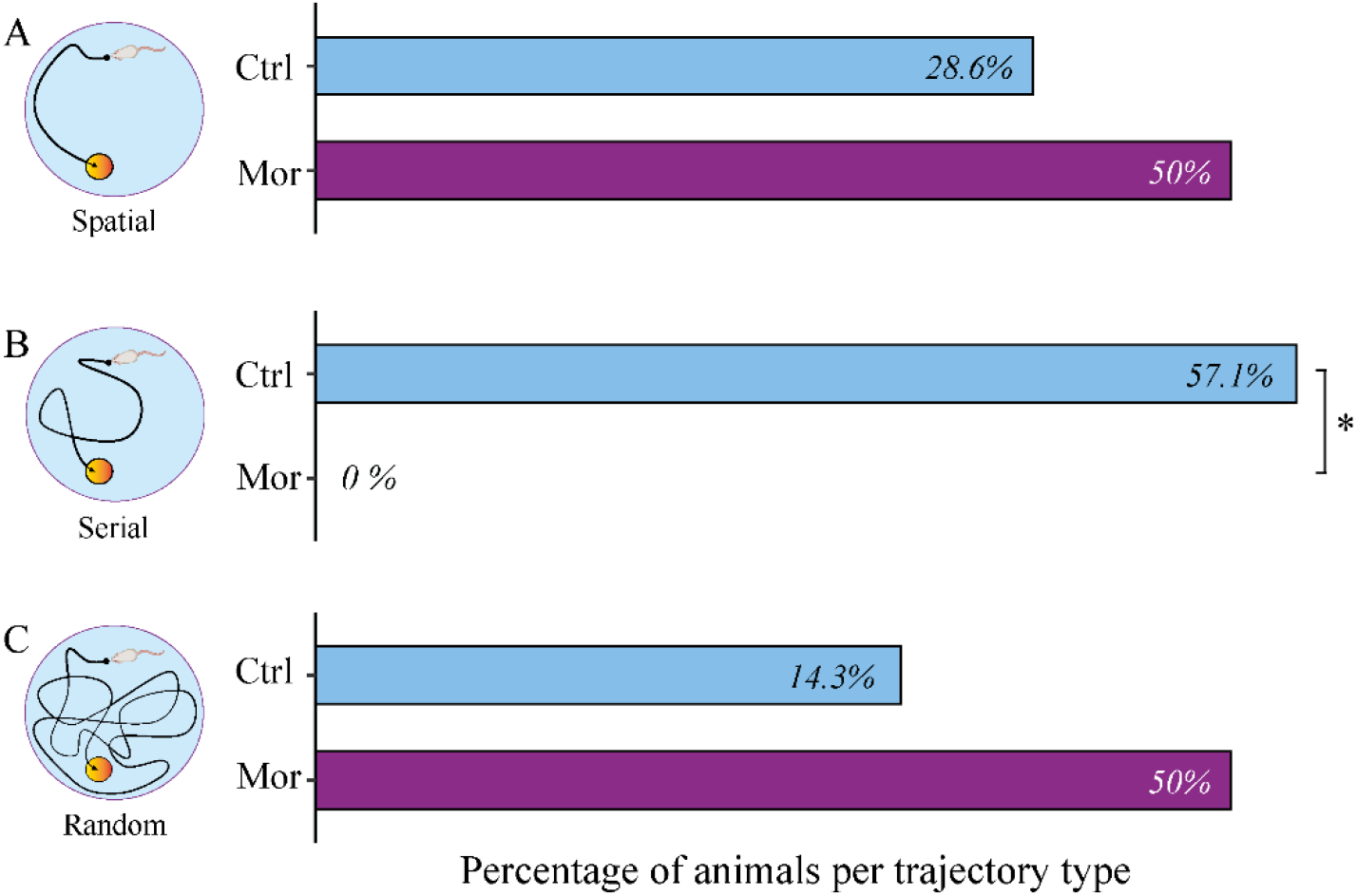
Distribution of swimming trajectory strategies during the MWM probe test. Representative schematic diagrams illustrate the three swimming strategies. **A.**Spatial trajectories, defined as direct paths toward the platform area. **B**. Serial trajectories, defined as systematic exploration of some quadrants without traversing the entire maze to reach the platform area. **C**. Random trajectories, defined as non-directed swimming patterns characterized by exploration of multiple quadrants without a specific goal. Bars represent the percentage of animals in each group exhibiting each trajectory type. *p < 0.05, Fisher’s exact test.

### Morphine self-administration induces longitudinal brain volume remodeling, including lobule-specific cerebellar changes

Following behavioral characterization, longitudinal volumetric analyses were performed to assess brain structural changes. As shown in Fig. 5A, morphine-exposed rats produced a progressive and linear volume enlargement in the cingulate cortex (Cg1–2), primary somatosensory cortex (S1), auditory cortex (Au), visual cortex (V1), dorsal striatum (CPu), dentate gyrus (DG), posteromedial cortical amygdaloid nucleus (PMCo), and the cerebellum, including the 4^th^ cerebellar lobule (4Cb), flocculus (Fl), and lateral Crus1 of the ansiform lobule (L-Crus1). In contrast, volume reductions emerged in frontal cortex area 3 (Fr3), the insular cortex (Ins), nucleus accumbens core (AcBC), septofimbrial nucleus (Sfi), subiculum transition area (S), deep white layer of the superior colliculus (DpWh), microcellular tegmental nucleus (MiTg), paraflocculus (PFl), and medial Crus1 of the ansiform lobule (M-Crus1). Representative trajectories from 4Cb (Fig. 5B), L-Crus1 (Fig. 5C), and M-Crus1 (Fig. 5D) depict group-wise volume trajectories across the self-administration period relative to control animals. Morphine self-administration was associated with time-dependent, non-linear patterns of volumetric remodeling (Fig. 5E). Inverted U shape volume changes were observed in the ventral orbital cortex (VO), CPu, corpus callosum (CC), thalamic nucleus (Th), Au, pedunculopontine tegmental nucleus (PTg), and retrosplenial granular cortex (RSG), whereas U shape volume changes were found in the secondary motor cortex (M2), nucleus accumbens shell (AcBS), Ins, S1, medial mammillary nucleus, lateral part (ML), amygdalopiriform transition area (APir), S, presubiculum (PrS), and gigantocellular reticular nucleus, ventral part (GiV). Within the cerebellum, non-linear modeling differentiated lobule-specific temporal profiles. Crus1 exhibited an initial volume decrease followed by an increase (Fig. 5F). The 8^th^ cerebellar lobule (8Cb) showed an increment that stabilized over time, whereas the 7^th^ cerebellar lobule (7Cb) displayed a transient reduction followed by partial normalization (Fig. 5G-H). The complete lists of regions are provided in Supplementary Tables S3 and S4.

**Figure 5.**
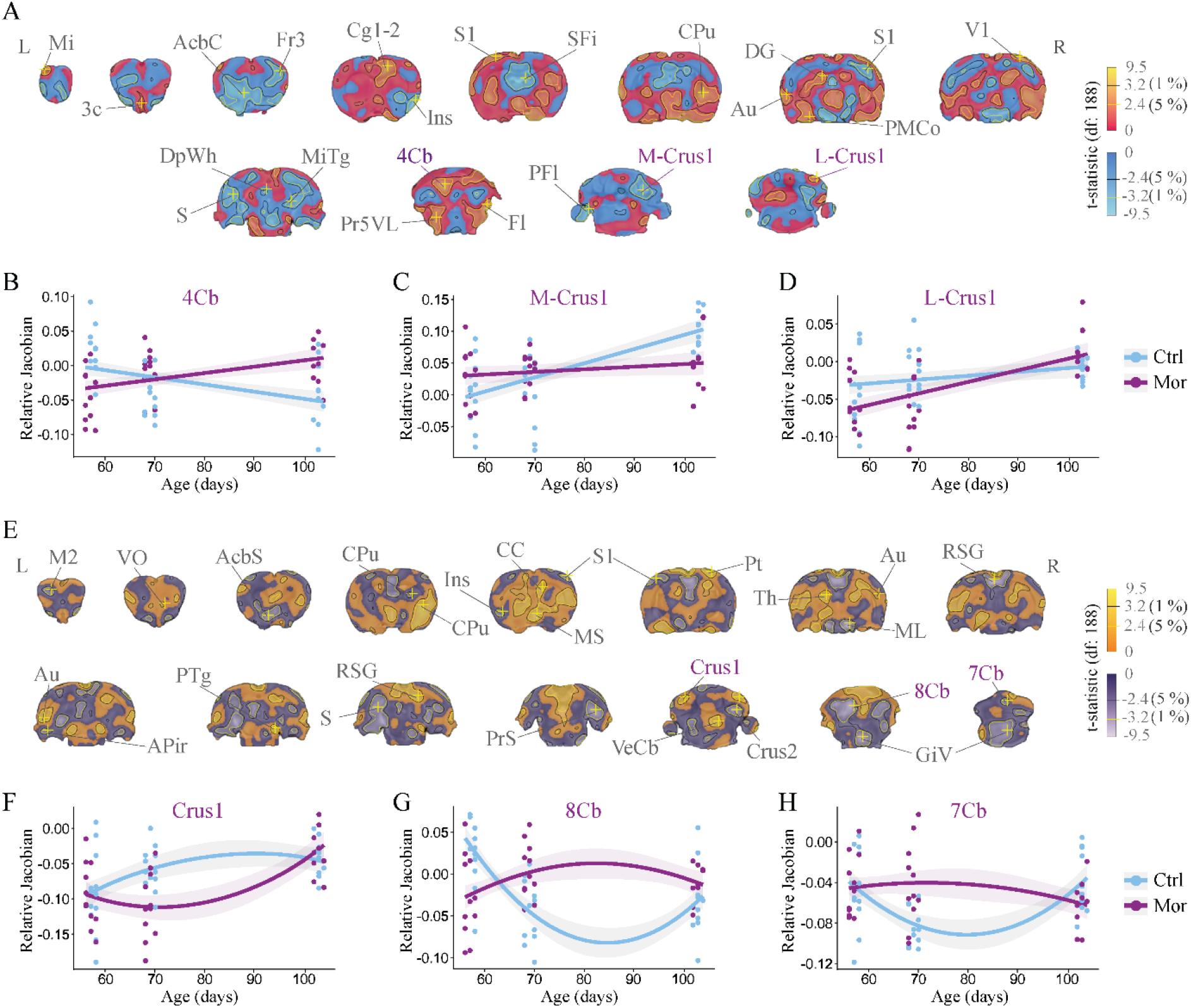
Longitudinal deformation-based morphometry analysis of brain volume across morphine self-administration. **A.**Coronal sections from rostral to caudal showing voxel-wise longitudinal linear effects on brain volume (Group × Age mixed-effects model). **B-D**. Linear trajectories (Relative Jacobians) across postnatal age for selected regions (4Cb, M-Crus1, L-Crus1), derived from MRI at T1 (P57-58), T2 (P69-70), and T3 (P104-105). **E**. Coronal sections showing voxel-wise quadratic longitudinal effects on brain volume (Group × Age^2^ mixed-effects model). **F-H**. Plots illustrating model-estimated quadratic trajectories for Crus1 (F), 8Cb (G), and 7Cb (H). Points represent individual animals; lines indicate model fits; shaded areas denote 95% confidence intervals. Statistical maps were obtained from voxel-wise LMMs (df = 188). Colors indicate t-statistic sign (cool: negative; warm: positive). Voxel-wise significance was controlled using FDR (q < 0.05), with q < 0.01 outlined in yellow; the |t| = 2.4 contour (black) corresponds to the FDR threshold. Coronal slices are labeled according to the Paxinos and Watson stereotaxic atlas; abbreviations are listed in Supplementary Table S2. Ctrl, control; Mor, morphine. L, left hemisphere; R, right hemisphere; P, postnatal day.

### Cerebellar functional connectivity is reorganized following morphine self-administration

We examined resting state functional connectivity (FC) using seed-based resting-state fMRI. Seeds were defined from cerebellar clusters showing robust volumetric effects (7/8Cb and Crus1/cbw clusters; Fig. 5F-H)). Specifically, we observed decreased connectivity of lobule 7/8Cb with S1, Ins, CPu, hippocampal field CA2, and the thalamus (ventral lateral nucleus; VL). We also found increased FC of 7/8Cb with the primary motor cortex (M1), the corpus callosum (forceps minor; fmi), and additional cerebellar regions, including lobules 3 and 4 (3/4Cb) and lobules 8, 9, and 10 (8/9/10Cb) (Fig. 6B, C). The Crus1/cbw seed region (Fig. 6D) exhibited reduced FC with auditory cortex (ventral area; AuV), cingulate cortex (area 2; Cg2), prelimbic cortex (PrL), hippocampal field CA1, CPu, VL, and additional cerebellar regions, including lobules 3 and 4 (3/4Cb) and lobules 8 and 9 (8/9Cb). We also found increased FC with entorhinal cortex (Ent), secondary visual cortex (V2), M1, the olfactory bulb (granular layer; GrO), and S1 (Fig. 6E, F). Both seed regions (7/8Cb and Crus1/cbw) also showed increased and decreased FC with the reticular nucleus (mRt) respectively. Complementary analyses using the hippocampus CA2, dentate gyrus (DG), and striatum as additional seeds revealed distributed changes in FC across cortical, hippocampal, thalamic, striatal, and brainstem regions (Supplementary Fig. S1), several of which overlapped with those identified in the cerebellar connectivity maps.

**Figure 6.**
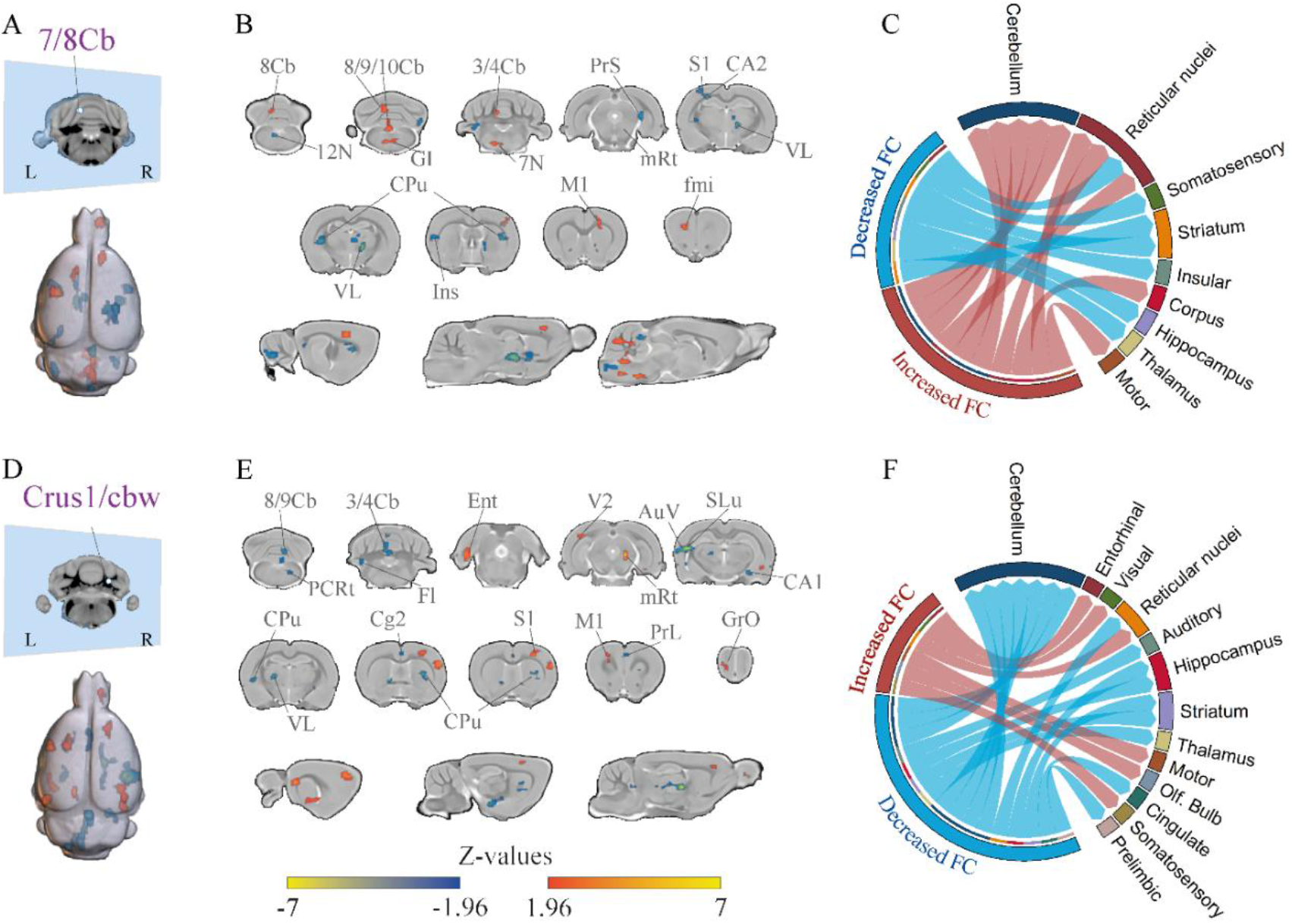
Cerebellar seed-based functional connectivity associated with morphine self-administration. **A.**Cerebellar seed including lobules 7/8. **B**. Voxel-wise connectivity maps for the 7/8 cerebellar seed, showing regions with decreased (blue) and increased (red) coupling in morphine-exposed animals. **C**. Circular plot summarizing connectivity changes between lobules 7/8 and cortical and subcortical systems. **D**. Cerebellar seed including Crus1 and adjacent cerebellar white matter (cbw). **E**. Voxel-wise connectivity maps for the Crus1/cbw seed. **F**. Connectivity summary for the Crus1/cbw seed across distributed brain systems. Statistical maps were thresholded at |Z| ≥ 1.96 and cluster-corrected using nearest-neighbor voxel contiguity (NN = 1). Abbreviations of brain regions are listed in Supplementary Table S2. FC, functional connectivity; L, left; R, right.

### Motivational drive is linked to cerebellar and addiction-related brain network patterns

To examine multivariate associations between brain structure and behavioral performance, we conducted a partial least squares (PLS) analysis relating brain volume to behavioral measures. Permutation testing indicated that the first latent variable (LV1) accounted for approximately 61% of the shared covariance; however, this effect did not reach statistical significance (p = 0.057; Fig. 7A). Despite this, LV1 showed a group-dependent pattern in both behavioral and brain scores, with morphine-exposed animals displaying higher expression compared to controls (behavioral scores: p = 0.002; brain scores: p = 0.04; Fig. 7B). Inspection of behavioral loadings revealed that motivational measures contributed most strongly to LV1, as indicated by bootstrap ratios whose confidence intervals excluded zero (Fig. 7C), whereas other behavioral variables showed fewer stable contributions. The corresponding brain pattern included distributed regions spanning cerebellar (Crus1, Crus2, 6Cb), insular, striatal, hippocampal, and amygdaloid areas, as well as thalamic and brainstem nuclei (Fig. 7D). This spatial pattern overlapped with regions identified in volumetric analyses. Together, these results suggest a potential association between motivational performance and distributed brain structural variation.

**Figure 7.**
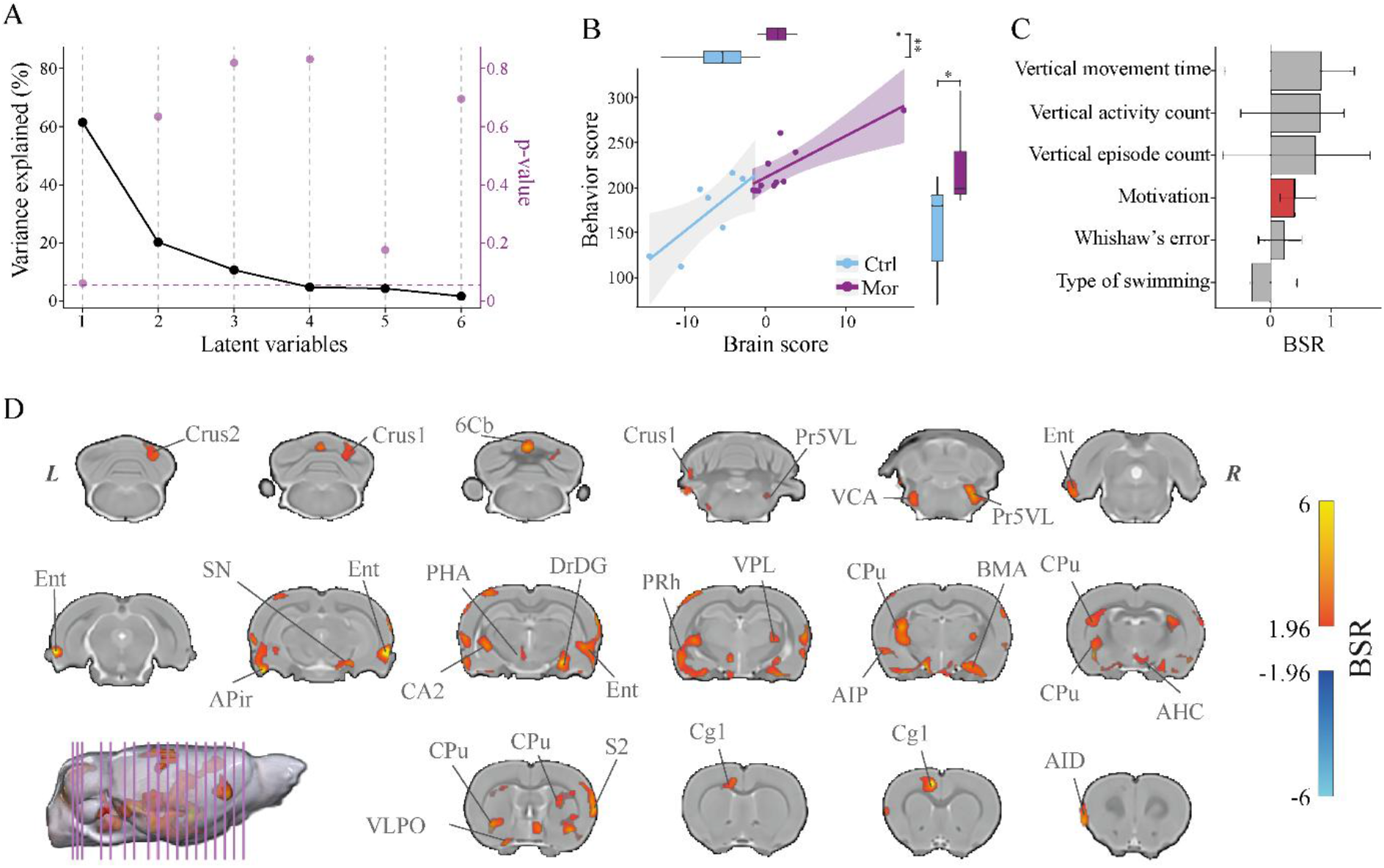
Relationship between brain structural measures and behavioral variables assessed using Partial Least Squares (PLS) analysis. **A.**Percentage of variance explained by each latent variable (black line), with corresponding permutation-based p-values shown in purple; the dashed horizontal line indicates the p = 0.05 significance threshold. **B**. Distribution of individual brain and behavioral scores for LV1 by group (Mor vs Ctrl). *q < 0.05, **q < 0.01, ***q < 0.001. **C** Behavioral loadings contributing to LV1. Vertical exploration includes rearing count, episodes, and total time; motivation is derived from lever pressing; Whishaw’s error and swimming strategy were obtained from the Morris water maze. Colors indicate significant (red) and non-significant (gray) contributions. **D**. Brain maps showing voxel-wise BSR values for LV1 (cool: negative; warm: positive; |BSR| ≥ 1.96). Abbreviations are listed in Supplementary Table S2.

## 4. Discussion

Opioid addiction has traditionally been framed as a dysfunction of reward circuitry (Koob and Volkow 2016). However, emerging evidence suggests that cerebellar networks contribute to motivated actions as well. Using behavioral, structural MRI, functional connectivity, and multivariate analyses, we showed that morphine self-administration produces a dissociation between preserved motivational drive and impaired organization of goal-directed behavior, supported by altered integration between cerebellar and cerebral networks.

Motivational engagement during morphine self-administration first became evident during the FR1 acquisition phase, as reflected by the progressive increase in active lever presses. The early establishment of this response is consistent with the strong reinforcing efficacy of opioids when response costs are low (Swain et al. 2021). The transition to the PR schedule further amplified this effect, preserving sustained responding despite increasing response costs, a feature of enhanced motivational drive (Ahmed and Koob 1998; Everitt and Robbins 2016). Motivational scores confirmed that morphine-exposed animals were strongly motivated to obtain the drug. This profile reflects addiction-like phenotypes in which progressive-ratio lever pressing serves as an index of motivational drive for the drug reward (Bock et al. 2013; Belin et al. 2013).

Behavioral analyses indicated that morphine self-administration at this dose produced selective rather than generalized effects. Enhanced exploratory activity emerged following stabilization of morphine intake. Previous evidence indicates that rearing contributes to the formation of spatial representations through sampling of distal environmental cues (Shan et al. 2023) and increases after exposure to μ-opioid receptor agonists such as oxycodone (De Almeida et al. 2024). In contrast, anxiety-like behavior and recognition memory remained largely unaffected, while MWM performance indicated preserved spatial learning, consistent with that reported in addiction (Everitt and Robbins 2016; Heinsbroek et al. 2021). However, morphine-exposed rats displayed impairments in the organization of goal-directed behavior. Specifically, these animals exhibited increased Whishaw’s error (Whishaw and Mittleman 1986), indicating reduced navigational precision despite preserved spatial memory. The coexistence of increased exploratory rearing and elevated Whishaw’s error suggests that morphine self-administration did not reduce environmental engagement but disrupted the translation of spatial information into organized action sequences, potentially involving cerebellar circuits interacting with basal ganglia, hippocampal, and thalamic networks (Rochefort et al. 2011; Bostan and Strick 2010). Swimming strategy analysis further supported this interpretation, as control animals relied on a balanced combination of spatial and serial trajectories, whereas morphine exposure abolished serial strategies and increased random navigation patterns. Because serial strategies depend on efficient movement sequencing (Vorhees and Williams 2006), their absence indicates disrupted action organization during navigation. Within addiction contexts, such dissociations suggest that drug use may preserve goal pursuit while degrading the efficiency of action execution (Everitt and Robbins 2016).

The morphine-induced behavioral phenotype observed in this study aligns with the role of the cerebellum in coordinating goal-directed behavior. Beyond motor coordination, the cerebellum contributes to action planning, sequencing, temporal prediction, and the automation of learned behavioral routines through distributed cerebello-cerebral networks involving prefrontal, limbic, and striatal regions (Buckner 2013; Rudolph et al. 2023; Manto et al. 2024; Therrien and Bastian 2015). Accordingly, morphine self-administration produced progressive cerebellar remodeling alongside volumetric changes across cortical, striatal, hippocampal, and thalamic regions. These effects may reflect, in part, the action of morphine on cerebellar circuits expressing μ-opioid receptors (Kinney and White 1991; Bekheet et al. 2010; Yang et al. 2022) and their integration with motor and prefrontal networks (Kelly and Strick 2003; Rudolph et al. 2023). Furthermore, cerebellar volume alterations followed lobule-specific linear and non-linear trajectories across Crus1, 7Cb, 8Cb, and cerebellar white matter, suggesting experience-dependent structural plasticity reported in prior studies (Lerch et al. 2017; Fjell et al. 2015). Extending our previous findings (Elizarrarás-Herrera et al. 2026), longer morphine exposure produced more robust cerebellar remodeling, reinforcing the involvement of cerebellar circuits in drug-seeking behavior.

To further examine the functional implications of cerebellar remodeling, resting-state fMRI analyses were performed using cerebellar seeds derived from clusters showing robust volumetric effects. This approach was guided by anatomical evidence demonstrating that the cerebellum participates in circuits linking specific lobules with cortical and subcortical regions (Middleton and Strick 2001; Kelly and Strick 2003). Cerebellar remodeling was associated with altered functional coupling across motivational and sensorimotor networks. Alterations in lobules 7/8 were characterized by reduced connectivity with insular, striatal, thalamic, and hippocampal regions, together with increased coupling to motor and intra-cerebellar territories. Similarly, changes involving Crus1 showed reduced interactions with the prefrontal cortex and increased connectivity with entorhinal and sensory association cortices. These regions participate in circuits involved in interoception, motivational salience, and action selection, and the observed connectivity pattern is consistent with circuit-level imbalance models proposed in addiction (Craig 2003; Naqvi and Bechara 2009; Zhang et al. 2025; Müller-Oehring et al. 2022). Complementary analyses using striatal, hippocampal, and interhemispheric seeds revealed partially overlapping connectivity patterns with those observed using cerebellar seeds. Together, these findings suggest that morphine-related alterations in cerebellar connectivity may disrupt the integration of motivational and sensorimotor signals required for the efficient organization of goal-directed actions during drug seeking.

Complementary to region-specific functional coupling, multivariate approaches have emphasized the importance of examining integrated brain-behavior relationships to capture neural contributions to behavior in addiction (Moulton et al. 2014; Moreno-Rius and Miquel 2017; Bostan et al. 2013). Our covariance analysis identified a dominant latent variable linking behavioral and structural measures. The observed pattern suggested a potential association between motivational drive for morphine-seeking and brain volume variation. The corresponding neuroanatomical profile included cerebellar and addiction-related regions, such as the insula, striatum, hippocampus, and amygdala, which are involved in interoception, incentive valuation, contextual processing, and affective salience, respectively (Koob and Volkow 2016; Naqvi and Bechara 2009), with cerebellar circuitry potentially contributing to the integration of these processes (Moulton et al. 2014). The multivariate result overlapped with regions identified in volumetric and functional connectivity analyses, indicating convergence across analytical approaches and supporting the presence of circuit-level adaptations. At the behavioral level, motivational measures showed the most stable contribution, consistent with the robust expression of drug-seeking behavior observed under the progressive-ratio schedule. Together, these findings suggest that motivational drive is the behavioral dimension most strongly associated with distributed structural variation across cerebellar and addiction-related circuits.

Limitations of this study should be considered when interpreting the findings. Only male Wistar rats were included, which may limit generalizability given potential sex differences in opioid-related behaviors. Although the longitudinal design allowed parallel assessment of behavior and brain changes, these analyses remain correlational and do not establish causality. The extensive behavioral battery may have contributed to a modest reduction in morphine intake toward the end of the protocol. Additionally, behavioral assays probing cerebellar integrative functions remain limited.

## 5. Conclusion

In conclusion, morphine self-administration produces a dissociation between preserved drug-seeking motivation and impaired organization of goal-directed actions. Cerebellar remodeling and altered cerebello-cerebral coupling appear to contribute to the integration of motivational signals into organized action sequences during drug seeking. These findings extend reward-centered views of opioid addiction by highlighting the cerebellum as an important component of circuit-based models shaping motivated behavior.

## Acknowledgements

This work was partially derived from the master’s thesis of Mariana Stefania Serrano-Ramírez at the Institute of Neurobiology, UNAM. Technical assistance was provided by the University Animal Facility Laboratory of the INB, particularly José Martín García Servín, María A. Carbajo Mata, and Alejandra Castilla León. Soledad Mendoza Trejo from Laboratory D12 provided technical support during the experimental procedures. Dr. Deisy Gasca Martínez from the Behavioral Analysis Unit. Computational support was provided by Ramón Martínez Olvera, Moisés Mendoza Baltzar, and María Eugenia Rosas Alatorre. Magnetic resonance imaging data were acquired at the National Laboratory for Magnetic Resonance Imaging (LANIREM) with assistance from Luis Concha, Juan José Ortíz Retana, and Leopoldo González Santos. Computational resources were provided by Mallar Chakravarty and Gabriel A. Devenyi at the Computational Brain Anatomy Laboratory (CoBrA Lab) (http://cobralab.ca/), CIC, Douglas Research Centre, Montreal, and Digital Research Alliance Canada (https://alliancecan.ca). Access to computational tools on the Trillium Compute Cluster was provided by Mallar Chakravarty, Director of the Computational Neuroanatomy Laboratory at the Douglas Research Centre (Montreal, Canada).

## Funding

This work was supported by the UNAM PAPIIT projects IN213924 and IA201622 (EAGV) and partially supported by postdoctoral fellowships from DGAPA-UNAM (629578; CJCA) and SECIHTI (3256252; CJCA).

## Author contribution

EAGV, CJCA, and MSSR conceptualized and designed the study. EAGV provided drugs, reagents, and equipment. CJCA, DMS, MSSR, and LATV performed the surgeries. CJCA, MSSR, JRT, and DMS conducted the behavioral experiments and acquired the MRI scans. JRT performed the structural and functional MRI analyses. JRT, CJCA, and LATV carried out the statistical analyses. The manuscript was drafted by CJCA, and all authors contributed to the revision and approved the final version.

## Notes

### Competing Interest Statement

The authors have declared no competing interest.

