## Supplementary material for "Cerebellar structural and functional alterations during morphine self-administration are associated with motivation and disrupted goal-directed actions in male Wistar rats"

#### Supplementary methods

##### Catheter implantation

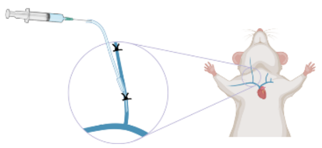

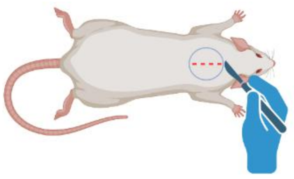

**Anesthesia**: Isoflurane (5% induction, ~1% maintenance). Adequate anesthesia was confirmed by absence of the pedal withdrawal reflex.
**Surgical preparation**: Ventral neck region shaved and disinfected with a superoxidized solution (Microdacyn®) followed by 70% ethanol.
**Catheter**: 22-gauge polyurethane catheter (C30PU-RJV1301).
**Insertion site**: Right jugular vein.
**Externalization**: Catheter tunneled subcutaneously to the interscapular region and connected to a non-magnetic vascular access button (VAB95BS; Instech Laboratories, Inc.).
**Wound closure**: Nylon sutures.
**Postoperative care**: Six-day recovery period with daily flushing (100 µl saline containing ticarcillin 66.67 mg/ml and heparin 20 IU/ml).
**Patency verification**: 100 µl ketamine (15 mg/ml) + midazolam (0.75 mg/ml) mixture producing rapid loss of the righting reflex.
**Maintenance**: Daily flushing with heparinized saline after each self-administration session.
**Monitoring**: Catheter sites inspected regularly for occlusion, infection, or other complications.

##### Operant training

**Operant chambers:** Med-Associates, ENV-018V.
**Shaping protocol:** Three-day shaping procedure to establish lever-press behavior (1).
  ***Day 1 (habituation):*** *Animals were placed in the chamber with the house light illuminated and one lever extended and allowed free exploration; food was removed overnight after the session.*
  ***Day 2 (successive approximations):*** *Lever-directed behaviors (orientation, approach, contact, and full lever pressing) were reinforced with food pellets; regular feeding was restored after the session.*
  ***Day 3 (FR1 shaping):*** *Rats responded under a fixed-ratio 1 schedule with food reinforcement available for 1 h.*
**Session timing:** All self-administration sessions began at 10:00 a.m. during the dark phase of the light–dark cycle.

##### Self-administration schedules and details

**Lever configuration:** Two-lever setup with counterbalanced active and inactive levers to establish drug-reinforced responding.
**FR1 phase:** 20 days; each active lever press delivered morphine (0.1 mg/kg/infusion; 100 µl over 5 s at 20 µl/s).
**PR phase:** Progressive-ratio 9-4 schedule (6 h/day, 25 days) with escalating response requirements (2).
**Cue light:** Yellow cue light located 5 cm above the active lever paired with each infusion.
**House light:** Remained illuminated during sessions.
**Timeout:** 30 s timeout after each infusion with no lights or active levers available.
**Infusion limit:** Maximum of 20 infusions per session.
**Recorded variables:** Active lever presses, inactive lever presses, and number of infusions.
**Motivation index:** Calculated from the final five PR sessions as the mean number of active lever presses emitted before the last infusion of each session (3,4). Higher values indicate greater motivational drive to obtain morphine.

#### Behavioral testing procedures

| Test | Apparatus and software | Procedure<br>(~2 h after self- administration/dark phase) | Behavioral measures |
|---|---|---|---|
| **Open Field Test (OFT)** | Open-field actimeter (Omnitech Electronics)<br>42 × 42 × 30 cm<br>16 × 16 infrared beam grid (2.5 cm spacing)<br>Software: Fusion SuperFlex v5.3 | Free exploration (10 min) | Rearing count<br>Rearing duration |
| **Novel Object Recognition (NOR)** | Acrylic arena<br>40 × 40 × 40 cm<br>Software: DeepLabCut (ResNet-50; RRID:SCR_021391) and DLCAnalyzer | Habituation (10 min)<br>Familiarization with two identical objects (10 min)<br>Test after 24 h with one novel object (10 min) | Discrimination ratio |
| **Elevated Plus Maze (EPM)** | Elevated plus maze<br>45 × 10 cm arms, 40 cm height<br>Software: DeepLabCut and DLCAnalyzer | Exploration (5 min)<br>Starting from central platform facing open arm | Anxiety index<br>Arm entries<br>Time in arms<br>Distance<br>Velocity |
| **Morris Water Maze (MWM)** | Circular pool<br>1.82 m diameter<br>Submerged platform (15 cm)<br>Software: Smart Video Tracking v3.0 | 4 training days (4 trials/day)<br>Probe test (60 s) without platform | Escape latency<br>Time in target quadrant<br>Entries<br>Directionality<br>Whishaw's error<br>Swimming strategy |

#### Magnetic Resonance Imaging

| | Parameter | Details |
|---|---|---|
| **MRI** | Location | National Laboratory for Magnetic Resonance Imaging (LANIREM) |
| | Scanner | 7 Tesla Bruker Pharmascan 70/16 US |
| | Coil | 2 × 2 rat head surface coil |
| | Acquisition software | Paravision v7.0 |
| **Anesthesia and monitoring** | Induction | 5% isoflurane |
| | Maintenance | ~1% isoflurane |
| | Gas mixture | 1:1 oxygen/air mixture |
| | Temperature | Circulating warm-water system set to 38 °C |
| | Physiological monitoring | Continuous monitoring of cardiac and respiratory signals |
| **Structural MRI acquisition** | Image type | High-resolution T2-weighted images |
| | Sequence | 3D FLASH |
| | TR | 30.76 ms |
| | TE | 5 ms |
| | Flip angle | 10° |
| | Field of view | 28.2 × 19 × 25.6 mm |
| | Voxel size | 160 μm isotropic |
| | Repetitions | 2 |
| | Primary slice direction | Sagittal |
| **Functional MRI acquisition** | Image type | Functional T2*-weighted images |
| | Sequence | GE-EPI |
| | TR | 1000 ms |
| | TE | 20 ms |
| | Flip angle | 60° |
| | Slice thickness | 1 mm |
| | Field of view | 30 × 30 mm |
| | Number of slices | 24 |
| | Volumes | 600 |
| **Deformation-Based Morphometry (DBM)** | Data conversion | Structural MRI data converted to BIDS format using the brkraw toolbox (v0.3.11) |
| | Preprocessing | Averaging repetitions to improve signal-to-noise ratio, intensity normalization, centering, denoising, and iterative co-registration |
| | Software | MINC-toolkit (v2; RRID:SCR_003519) and ANTs (RRID:SCR_004757) |
| | Quality control | Performed at all stages, with no exclusions |
| | Template workflow | Two-step workflow: individual sessions averaged to create subject-specific templates, followed by construction of a study-specific template serving as the common registration space |
| | Volume metric | Jacobian determinant maps reflecting local volume differences relative to the study average |
| | Atlas labeling | SIGMA rat brain atlas (v1.2.1) (5) |
| **Functional connectivity analysis** | Preprocessing pipeline | RABIES (v0.5) |
| | Preprocessing steps | Inhomogeneity correction, nonlinear registration to a common space, resampling to 0.3-mm³ voxels, motion correction |
| | Temporal filtering | High-pass (0.01 Hz) and low-pass (0.1 Hz) filtering |
| | Spatial smoothing | Gaussian smoothing (0.5 mm) |
| | Nuisance regression | Regression of white-matter, cerebrospinal-fluid, and motion-parameter signals |
| | Connectivity analysis | Seed-based functional connectivity |
| | Seed definition | Clusters with the highest t-statistic values derived from structural DBM results |
| | Seed masks | Binary inflated masks (radius = 0.4 mm) |
| | Signal extraction | Time-series signals extracted from seed regions |

#### Statistical analysis

All behavioral variables were inspected for distributional properties using the Shapiro-Wilk test. Self-administration measures (infusions, active and inactive lever presses, and body weight) were examined using linear mixed-effects models with rat as a random effect and group, session, and/or protocol (FR1/PR) as fixed effects, followed by Tukey-adjusted post hoc comparisons. OFT and EPM data were analyzed using two-way repeated-measures ANOVA (Group × Phase: T1/T2) with Bonferroni-adjusted post hoc tests. NOR performance was evaluated using Welch's independent-samples t-tests. MWM acquisition data were analyzed using linear mixed-effects models. Probe-test variables were analyzed using ordinary least squares (OLS) regression with Group as the predictor for continuous outcomes, and Poisson generalized linear models for count data. Swimming trajectory distributions were analyzed using Fisher's exact tests. A global 2 × 3 test (Group × Strategy) was followed, when appropriate, by Holm-adjusted 2 × 2 comparisons. Effect sizes were estimated as generalized eta squared ($\eta^2 g$) for ANOVA models, Hedges' g or Cohen's d for between-group comparisons, marginal $R^2$ for mixed-effects models and $R^2$ for linear models, incidence rate ratios (IRR) for Poisson models, and Cramér's V or phi (ϕ) for categorical analyses. To assess morphine-induced structural brain changes, voxel-wise linear mixed-effects models were applied (Local volume ~ Group × Age [or $Age^2$] + Batch + (1 | RID)), with multiple comparisons controlled using a 5% false discovery rate. Functional connectivity differences were tested using mixed-effects modeling (3dLMER; AFNI v23.2.04; RRID:SCR_005927) with the model Functional connectivity ~ Group × Age + Batch + (1 | RID). Brain volume-behavior associations were examined using Partial Least Squares (PLS) correlation, with 5000 permutations and split-half resampling used to assess statistical significance and reproducibility. Statistical significance for behavioral analyses was set at $p < 0.05$, with correction for multiple comparisons when applicable. Voxel-wise analyses were thresholded at $q < 0.05$ (FDR-corrected). All analyses were performed in R v4.2.3 (RRID:SCR_001905).

### Supplementary results

##### Supplementary Table S1. Morris Water Maze probe-test performance measures.

Summary of probe-test outcomes comparing morphine self-administering rats (Mor) and control animals (Ctrl). Distance is expressed in centimeters (cm), directionality and Whishaw's error in degrees, time in the target quadrant as proportion of total swim time, and entries as counts.

| Variable | Ctrl (Mean ± SD) | Mor (Mean ± SD) | Statistic | p-value | Effect size[a] |
|---|---|---|---|---|---|
| Mean Distance to Target | 71.76 ± 7.81 | 68.07 ± 12.39 | $t_{(15)} = -0.69$ | 0.751 | g = -0.32 |
| Mean Directionality to Target | 157.20 ± 149.45 | 180.10 ± 139.27 | $t_{(15)} = 0.32$ | 0.375 | g = 0.15 |
| Whishaw's Error | 93.11 ± 3.72 | 95.90 ± 2.25 | $t_{(15)} = 1.93$ | 0.036 | g = 0.90 |
| Time in Target Quadrant | 25.16 ± 18.09 | 21.68 ± 11.25 | $t_{(15)} = -0.49$ | 0.685 | g = -0.23 |
| Entries to Target Quadrant | 2.17 ± 1.47 | 2.90 ± 1.37 | z = 0.87 | 0.382 | IRR = 1.34 |

[a] *g: Hedges' g; IRR = incidence rate ratio.*

**Supplementary Table S2. Abbreviations of brain regions used in Figures 5-7.**
Brain divisions are organized according to major anatomical systems, and nomenclature follows the Paxinos and Watson rat brain atlas (6).

| Brain division | Abbreviation | Subregion |
|---|---|---|
| Cerebellum | 3Cb | 3rd Cerebellar lobule |
| | 4Cb | 4th Cerebellar lobule |
| | 6Cb | 6th Cerebellar lobule |
| | 7Cb | 7th Cerebellar lobule |
| | 8Cb | 8th Cerebellar lobule |
| | 9Cb | 9th Cerebellar lobule |
| | 10Cb | 10th Cerebellar lobule |
| | Crus1 | Crus I of the ansiform lobule, lateral (L), medial (M) |
| | Crus2 | Crus II of the ansiform lobule |
| | Fl | Flocculus |
| | PFl | Paraflocculus |
| | SLu | Simple lobule of the cerebellum |
| | VeCb | Vestibule cerebellar nucleus |
| | cbw | Cerebellar white matter |
| Cortex | 3c | Layer 3 of cortex |
| | Au | Auditory cortex |
| | AuV | Auditory cortex, ventral area |
| | Cg1 | Cingulate cortex, area 1 |
| | Cg2 | Cingulate cortex, area 2 |
| | Ent | Entorhinal cortex |
| | Fr3 | Frontal cortex area 3 |
| | Ins | Insular cortex |
| | M1 | Primary motor cortex |
| | M2 | Secondary motor cortex |
| | PrL | Prelimbic cortex |
| | PRh | Perirhinal cortex |
| | RSG | Retrosplenial granular cortex |
| | S1 | Primary somatosensory cortex |
| | S2 | Secondary somatosensory cortex |
| | V1 | Visual cortex |
| | V2 | Secondary visual cortex |
| | VO | Ventral orbital cortex |
| Hippocampal formation | CA1 | Hippocampal CA1 field |
| | CA2 | Hippocampal CA2 field |
| | DG | Dentate gyrus |
| | PrS | Presubiculum |
| | S | Subiculum, transition area |
| Basal ganglia | AcBC | Nucleus accumbens core |
| | AcBS | Nucleus accumbens shell |
| | CPu | Caudate-putamen (striatum) |
| | SN | Substantia nigra |
| Amygdala | AHC | Amygdalohippocampal area |
| | APir | Amygdalopiriform transition area |
| | BMA | Basomedial amygdaloid nucleus |
| | PMCo | Posteromedial cortical amygdaloid nucleus |
| | VCA | Ventral cortical amygdala |
| Thalamus/Hypothalamus | ML | Medial mammillary nucleus, lateral part |
| | PHA | Posterior hypothalamic area |
| | Th | Thalamic nucleus |
| | VL | Ventral lateral thalamic nucleus |
| | VLPO | Ventrolateral preoptic nucleus |
| | VPL | Ventral posterolateral thalamic nucleus |
| Brainstem | 7N | Facial nucleus |
| | 12N | Hypoglossal nucleus |
| | GI | Gigantocellular reticular nucleus |
| | GiV | Gigantocellular reticular nucleus, ventral part |
| | mRt | Mesencephalic reticular nucleus |
| | PCRt | Parvicellular reticular nucleus |
| | PTg | Pedunculopontine tegmental nucleus |
| | Pr5VL | Principal sensory nucleus of the trigeminal nerve, ventrolateral part |
| Other structures | CC | Corpus callosum |
| | DpWh | Deep white layer of the superior colliculus |
| | Mi | Mitral cell layer of the olfactory bulb |
| | MiTg | Microcellular tegmental nucleus |
| | Sfi | Septofimbrial nucleus |
| | fmi | Forceps minor of the corpus callosum |

**Supplementary table S3. Linear effects of morphine self-administration on brain volume.**

Brain regions showing significant linear volumetric effects identified using deformation-based morphometry (DBM). Sigma coordinates indicate peak voxel locations in SIGMA atlas space; main effect denotes the direction of change relative to controls.

| Brain region | Sigma coordinates (x, y, z) | Main effect | t-value |
|---|---|---|---|
| Agranular dysgranular insular cortex | 1.56, -0.74, -3.52 | Increase | 9.499 |
| Primary visual cortex | 3.69, -3.52, 4.83 | Increase | 7.461 |
| Molecular layer of the cerebellum | -2.22, -8.28, 2.21 | Increase | 7.443 |
| Striatum | 4.35, -1.39, -0.57 | Increase | 7.414 |
| Middle cerebellar peduncle | 4.02, -8.44, -0.9 | Increase | 7.341 |
| Primary auditory cortex | -7.63, -1.72, -0.41 | Increase | 7.223 |
| Cingular cortex | 1.23, 2.38, 3.36 | Increase | 6.639 |
| Brainstem | -3.53, -8.44, -2.7 | Increase | 5.581 |
| Dentate gyrus | -1.89, -2.71, 1.88 | Increase | 5.443 |
| Agranular dysgranular insular cortex | -2.87, -14.35, -1.56 | Increase | 5.371 |
| Entorhinal cortex | -4.35, -2.87, -3.69 | Increase | 5.325 |
| Primary somatosensory cortex forelimb | -3.69, 0.41, 4.5 | Increase | 5.304 |
| Agranular dysgranular insular cortex | 1.07, -5.82, -4.34 | Increase | 4.967 |
| Frontal association cortex | -1.56, 7.62, 3.52 | Increase | 4.723 |
| Granule cell level of the cerebellum | -7.3, -8.61, -1.23 | Increase | 4.653 |
| Molecular layer of the cerebellum | 3.2, -11.07, 3.03 | Increase | 4.602 |
| Primary somatosensory cortex upper lips | 5.66, 3.36, 0.9 | Increase | 4.588 |
| Olfactory bulb | -0.9, 12.21, 3.03 | Increase | 3.937 |
| Agranular dysgranular insular cortex | -4.68, 4.84, 3.52 | Increase | 3.698 |
| Agranular dysgranular insular cortex | -5.99, 1.07, -3.03 | Increase | 3.653 |
| Olfactory bulb | -0.41, 5.66, -1.56 | Increase | 3.413 |
| Agranular dysgranular insular cortex | -1.72, 2.71, -4.18 | Increase | 3.099 |
| Agranular dysgranular insular cortex | -0.74, -11.23, 4.5 | Decrease | -2.832 |
| Lateral parietal associative cortex | -4.68, -1.39, 3.19 | Decrease | -2.867 |
| Agranular dysgranular insular cortex | 6.81, -5.82, 1.39 | Decrease | -3.001 |
| Brainstem | -0.9, -8.12, -4.67 | Decrease | -3.211 |
| Agranular dysgranular insular cortex | -5.83, -9.59, 2.87 | Decrease | -3.738 |
| Deeper layers of the superior colliculus | -1.72, -5.66, 1.72 | Decrease | -4.172 |
| Brainstem | 1.39, -13.36, -3.85 | Decrease | -4.227 |
| Agranular dysgranular insular cortex | 2.71, 1.07, 5.16 | Decrease | -4.324 |
| Agranular dysgranular insular cortex | -2.38, 3.03, 5.16 | Decrease | -4.611 |
| Primary somatosensory cortex jaw | 4.02, 4.34, 2.87 | Decrease | -4.675 |
| Molecular layer of the cerebellum | 6.48, -10.9, -1.39 | Decrease | -4.88 |
| Agranular dysgranular insular cortex | 6.32, -5, -3.52 | Decrease | -4.899 |
| Brainstem | 1.72, -4.34, -0.08 | Decrease | -4.92 |
| Granule cell level of the cerebellum | 1.89, -10.25, 1.23 | Decrease | -5.846 |
| Olfactory bulb | 1.07, 10.9, 1.39 | Decrease | -5.956 |
| Corpus callosum and associated subcortical white matter | -5.99, -5, 1.23 | Decrease | -6.058 |
| Parietal cortex posterorostral | 4.68, -2.54, 3.19 | Decrease | -6.415 |
| Ventral hippocampal commissure | 0.41, 0.25, 1.23 | Decrease | -6.753 |
| Molecular layer of the cerebellum | -3.04, -12.87, 0.25 | Decrease | -7.329 |
| Prelimbic cortex | -1.56, 4.34, -0.08 | Decrease | -7.448 |
| Entorhinal cortex | 5.33, 2.54, -1.23 | Decrease | -7.542 |
| Granule cell level of the cerebellum | -5.5, -9.92, -1.39 | Decrease | -7.805 |
| Agranular dysgranular insular cortex | -1.56, -2.87, -3.52 | Decrease | -8.034 |

**Supplementary Table S4. Non-linear effects of morphine self-administration on brain volume.**

Brain regions showing significant non-linear volumetric effects identified using deformation-based morphometry. Sigma coordinates indicate peak voxel locations in SIGMA atlas space; main effect denotes the direction of change relative to controls.

| Brain region | Sigma coordinates (x, y, z) | Main effect | t-value |
|---|---|---|---|
| Molecular Layer of the Cerebellum | 0.08, -9.76, -0.57 | Increase | 8.414 |
| Striatum | 4.35, 0.25, -0.9 | Increase | 7.016 |
| Basal Forebrain Region | 0.25, 1.56, -1.88 | Increase | 7.01 |
| Lateral Parietal Associative Cortex | 2.87, -2.05, 5 | Increase | 6.865 |
| Retrosplenial Dysgranular Cortex | -1.23, -2.05, 5.32 | Increase | 6.708 |
| Molecular Layer of the Cerebellum | -5.33, -8.61, -1.56 | Increase | 6.292 |
| Parasubiculum | 1.56, -5.33,2.87 | Increase | 6.282 |
| Substantia Nigra | 2.05, -4.84, -2.05 | Increase | 6.106 |
| Agranular Dysgranular Insular Cortex | 3.04, -1.39, -4.67 | Increase | 6.085 |
| Agranular Dysgranular Insular Cortex | -4.02, -1.39, -4.5 | Increase | 6.033 |
| Corpus Callosum and Associated Subcortical White Matter | 0.74, 1.89, 2.21 | Increase | 5.803 |
| Thalamic Nucleus | -2.54, -3.2, 1.23 | Increase | 5.764 |
| Agranular Dysgranular Insular Cortex | -1.89, -7.3, 3.52 | Increase | 5.67 |
| Molecular Layer of the Cerebellum | 3.36, -10.9, 2.7 | Increase | 5.459 |
| Agranular Dysgranular Insular Cortex | -6.15, 2.54, 2.87 | Increase | 5.431 |
| Entorhinal Cortex | -7.47, -4.34, -0.41 | Increase | 5.346 |
| Agranular Dysgranular Insular Cortex | 7.3, -5.98, -0.57 | Increase | 5.102 |
| Granule Cell Level of the Cerebellum | 6.32, -9.76, -2.05 | Increase | 4.78 |
| Molecular Layer of the Cerebellum | -4.84, -10.41, 2.38 | Increase | 4.148 |
| Inferior Cerebellar Peduncle/Spinal Trigeminal Tract | -4.18, -12.54, -3.03 | Increase | 3.78 |
| Olfactory Bulb | -0.57, 12.21, 3.69 | Increase | 3.128 |
| Olfactory Bulb | -1.07, 5.16, 0.25 | Increase | 3.091 |
| Primary Auditory Cortex | 5.5, -3.2, 1.72 | Increase | 2.951 |
| Agranular Dysgranular Insular Cortex | 0.25, 6.31, 4.18 | Increase | 2.806 |
| Agranular Dysgranular Insular Cortex | -7.3, -0.25, -1.56 | Increase | 2.649 |
| Agranular Dysgranular Insular Cortex | 2.71, -9.59, -4.67 | Increase | 2.529 |
| Agranular Dysgranular Insular Cortex | 1.23, 9.1, -0.25 | Decrease | -2.822 |
| Agranular Dysgranular Insular Cortex | -2.54, 11.23, 2.21 | Decrease | -3.465 |
| Lateral Entorhinal Cortex Internal part | -6.97, -4.51, -2.37 | Decrease | -3.917 |
| Primary Motor Cortex | -2.22, 6.31, 2.7 | Decrease | -4.347 |
| Agranular Dysgranular Insular Cortex | 6.15, -1.56, -3.52 | Decrease | -4.673 |
| Entorhinal Cortex | -5.66, 1.56, -1.72 | Decrease | -4.725 |
| Descending Corticofugal Pathways and Globus Pallidum | 2.05, 0.9, 1.23 | Decrease | -4.745 |
| Agranular Dysgranular Insular Cortex | -6.32, -10.74, -2.87 | Decrease | -4.835 |
| Agranular Dysgranular Insular Cortex | 2.54, 3.03, -4.01 | Decrease | -4.857 |
| Agranular Insular Cortex | 4.68, 4.51, 0.25 | Decrease | -4.926 |
| Agranular Dysgranular Insular Cortex | -6.48, -10.57, 1.72 | Decrease | -5.025 |
| Primary Somatosensory Cortex Dysgranular | 5.01, 1.56, 3.85 | Decrease | -5.202 |
| Molecular Layer of the Cerebellum | -0.41, -12.21, 1.88 | Decrease | -5.249 |
| Agranular Dysgranular Insular Cortex | 1.89, 8.28, 3.69 | Decrease | -5.255 |
| Primary Somatosensory Cortex Barrel field | -5.99, -1.72, 4.01 | Decrease | -5.689 |
| Basal Forebrain Region | -1.23, 3.85, -1.88 | Decrease | -5.795 |
| Molecular Layer of the Cerebellum | 3.2, -10.9, 0.57 | Decrease | -6.24 |
| Perirhinal Area 36 | 4.35, -6.64,0.74 | Decrease | -6.557 |
| Brainstem | -0.57, -8.44, -3.36 | Decrease | -6.674 |
| Hypothalamic Region | 1.07, -3.2, -3.03 | Decrease | -6.76 |
| Subiculum | -5.33, -5.66, 1.23 | Decrease | -6.79 |
| Periventricular Grey | -0.25, -13.03, -3.03 | Decrease | -6.808 |
| Molecular Layer of the Cerebellum | -2.22, -8.44, 1.72 | Decrease | -7.788 |
| Retrosplenial Granular Cortex Part B | -0.9, -3.85, 3.85 | Decrease | -8.938 |

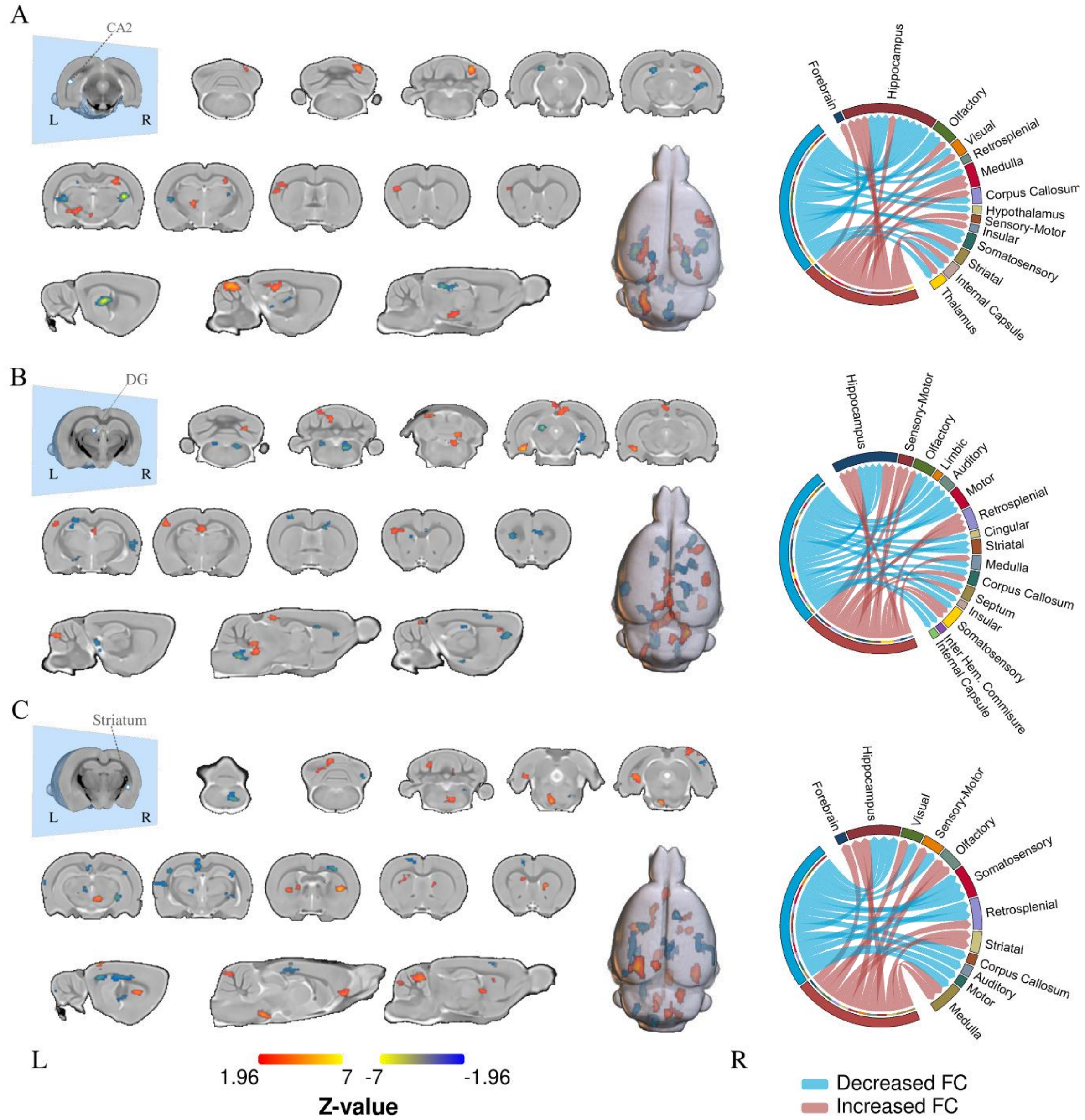


**Supplementary Figure S1.** Seed-based functional connectivity maps using additional anatomical seeds. **A**. Hippocampal CA2 field (CA2), **B.** Dentate gyrus (DG), and **C**. Striatum. Brain maps show regions with increased (warm colors) or decreased (cool colors) functional connectivity. Z-values correspond to the color scale shown below. Chord diagrams summarize connectivity changes across major brain systems. Statistical maps were thresholded at $|Z| \geq 1.96$ and cluster-corrected using nearest-neighbor voxel contiguity (NN = 1).
